# Leveraging Functional Annotations in Genetic Risk Prediction for Human Complex Diseases

**DOI:** 10.1101/058768

**Authors:** Yiming Hu, Qiongshi Lu, Ryan Powles, Xinwei Yao, Fang Fang, Xinran Xu, Hongyu Zhao

## Abstract

Genome wide association studies have identified numerous regions in the genome associated with hundreds of human diseases. Building accurate genetic risk prediction models from these data will have great impacts on disease prevention and treatment strategies. However, prediction accuracy remains moderate for most diseases, which is largely due to the challenges in identifying all the disease-associated variants and accurately estimating their effect sizes. We introduce AnnoPred, a principled framework that incorporates diverse functional annotation data to improve risk prediction accuracy, and demonstrate its performance on multiple human complex diseases.

## Main

Achieving accurate disease risk prediction using genetic information is a major goal in human genetics research and precision medicine. Accurate prediction models will have great impacts on disease prevention and early treatment strategies [1]. Advancements in high-throughput genotyping technologies and imputation techniques have greatly accelerated discoveries in genome-wide association studies (GWAS) [2]. Various approaches that utilize genome-wide data in genetic risk prediction have been proposed, including machine-learning models trained on individual-level genotype and phenotype data [3–8], and polygenic risk scores (PRS) estimated using GWAS summary statistics [9, 10]. Despite the potential information loss in summary data, PRS-based approaches have been widely adopted in practice since the summary statistics for large-scale association studies are often easily accessible [11, 12]. However, prediction accuracies for most complex diseases remain moderate, which is largely due to the challenges in both identifying all the functionally relevant variants and accurately estimating their effect sizes in the presence of linkage disequilibrium (LD) [13].

Explicit modeling and incorporation of external information, e.g. pleiotropy [7, 8] and LD [10], has been shown to effectively improve risk prediction accuracy. Recent advancements in integrative genomic functional annotation, coupled with the rich collection of summary statistics from GWAS, have enabled increase of statistical power in several different settings [14, 15]. To our knowledge, the impact of functional annotations on performance of genetic risk prediction has not been systematically studied.

Here, we introduce AnnoPred (available at https://github.com/yiminghu/AnnoPred), a principled framework that integrates GWAS summary statistics with various types of annotation data to improve risk prediction accuracy. We compare AnnoPred with state-of-the-art PRS-based approaches and demonstrate its consistent improvement in risk prediction performance using both simulations and real data of multiple human complex diseases.

AnnoPred risk prediction framework has three main stages (**Methods**). First, we estimate GWAS signal enrichment in 61 different annotation categories, including functional genome predicted by GenoCanyon scores [14], GenoSkyline tissue-specific functionality scores of 7 tissue types [15], and 53 baseline annotations [16]. Second, we propose an empirical prior of SNP effect size based on annotation assignment and signal enrichment. In general, SNPs located in annotation categories that are highly enriched for GWAS signals receive a higher effect size prior. Finally, the empirical prior is adopted in a Bayesian framework in which marginal summary statistics and LD matrix are jointly modeled to infer the posterior effect size of each SNP. AnnoPred PRS is defined by

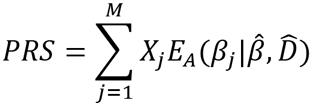

where *X_j_* and *β_j_* are the standardized genotype and effect size of the j^th^ SNP, respectively, 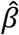 is the marginal estimate of *β*, 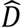 is the sample LD matrix, and *E_A_* denotes the posterior expectation under an empirical prior based on annotation assignment for all SNPs in the dataset (**Methods**).

We first performed simulations to demonstrate AnnoPred’s ability to improve risk prediction accuracy. We compared AnnoPred with four popular PRS approaches (**Methods**), i.e. PRS based on genome-wide significant SNPs (PRS_sig_), PRS based on all SNPs in the dataset (PRS_all_), PRS based on tuned cutoffs for p-values and LD pruning (PRS_P+T_), and recently proposed LDpred [10]. Mean correlations between simulated and predicted traits were calculated from 100 replicates under different simulation settings (Methods). AnnoPred showed the best prediction performance in all settings (**Table 1**). In general, performance of PRS_sig_, PRS_P+T_, LDpred, and AnnoPred all improved under a sparser genetic model and higher trait heritability. PRS_all_ showed comparable performance between sparse and polygenic models but its prediction accuracy was consistently worse than other methods. Sample size in the training set was also crucial for risk prediction accuracy. Doubling the training samples led to about 1.5-fold increase in AnnoPred’s performance under different settings in our simulations.

**Table 1.** Mean correlation between simulated and predicted traits calculated from 100 replicates under different simulation settings. The highest mean correlations are highlighted in boldface. Standard deviations are shown in parentheses.

| Training samples | Heritability | #Causal | PRS<sub>sig</sub> | PRS<sub>all</sub> | PRS<sub>P+T</sub> | LDpred | AnnoPred |
| --- | --- | --- | --- | --- | --- | --- | --- |
| Half | 0.25 | 300 | 0.149(.028) | 0.08(.021) | 0.25(.028) | 0.279(.025) | **0.286**(.024) |
|  |  | 3000 | NA* | 0.082(.016) | 0.073(.020) | 0.087(.019) | **0.096**(.020) |
|  | 0.5 | 300 | 0.304(.04) | 0.16(.022) | 0.48(.026) | 0.502(.033) | **0.512**(.026) |
|  |  | 3000 | NA* | 0.157(.019) | 0.157(.024) | 0.195(.021) | **0.209**(.019) |
| Full | 0.25 | 300 | 0.217(.031) | 0.11(.02) | 0.332(.023) | 0.35(.033) | **0.358**(.022) |
|  |  | 3000 | NA* | 0.11(.014) | 0.107(.018) | 0.136(.017) | **0.145**(.017) |
|  | 0.5 | 300 | 0.373(.036) | 0.213(.023) | 0.548(.024) | 0.557(.047) | **0.566**(.034) |
|  |  | 3000 | 0.078(.023) | 0.21(.019) | 0.243(.021) | 0.309(.021) | **0.324**(.019) |

\* NA means no SNP achieves genome-wide significance level (5e-8).

To further illustrate the improvement in risk prediction performance, we applied AnnoPred to five human complex diseases -- Crohn’s disease (CD), breast cancer (BC), rheumatoid arthritis (RA), type-II diabetes (T2D), and celiac disease (CEL). We first estimated GWAS signal enrichment in different annotation categories (**Methods**). Enrichment pattern varies greatly across diseases (**Figure 1A**; **Supplementary Table 1**), reflecting the genetic basis of these complex phenotypes. Functional genome predicted by GenoCanyon scores was consistently and significantly enriched for all five diseases. Blood was strongly enriched for three immune diseases, namely CD (P=8.9×10^−12^), CEL (P=7.0×10^−15^), and RA (P=9.9×10^−6^), while gastrointestinal (GI) tract was enriched in CD (P=2.6×10^−5^) and CEL (P=1.4×10^−4^), both of which have a known GI component. For BC, epithelium (P=7.4×10^−4^), GI (P=5.9×10^−3^), and muscle (P=6.1×10^−3^) were significantly enriched. Next, we evaluated the effectiveness of proposed empirical effect size prior in three diseases (i.e. CD, CEL, and RA) with well-powered testing cohorts (N>2,000). Interestingly, despite the highly variable enrichment results in training datasets, integrative effect size prior could effectively identify SNPs with large effect sizes and consistent effect directions in independent validation cohorts (**Figures 1B and 1C**).

**Figure 1.**
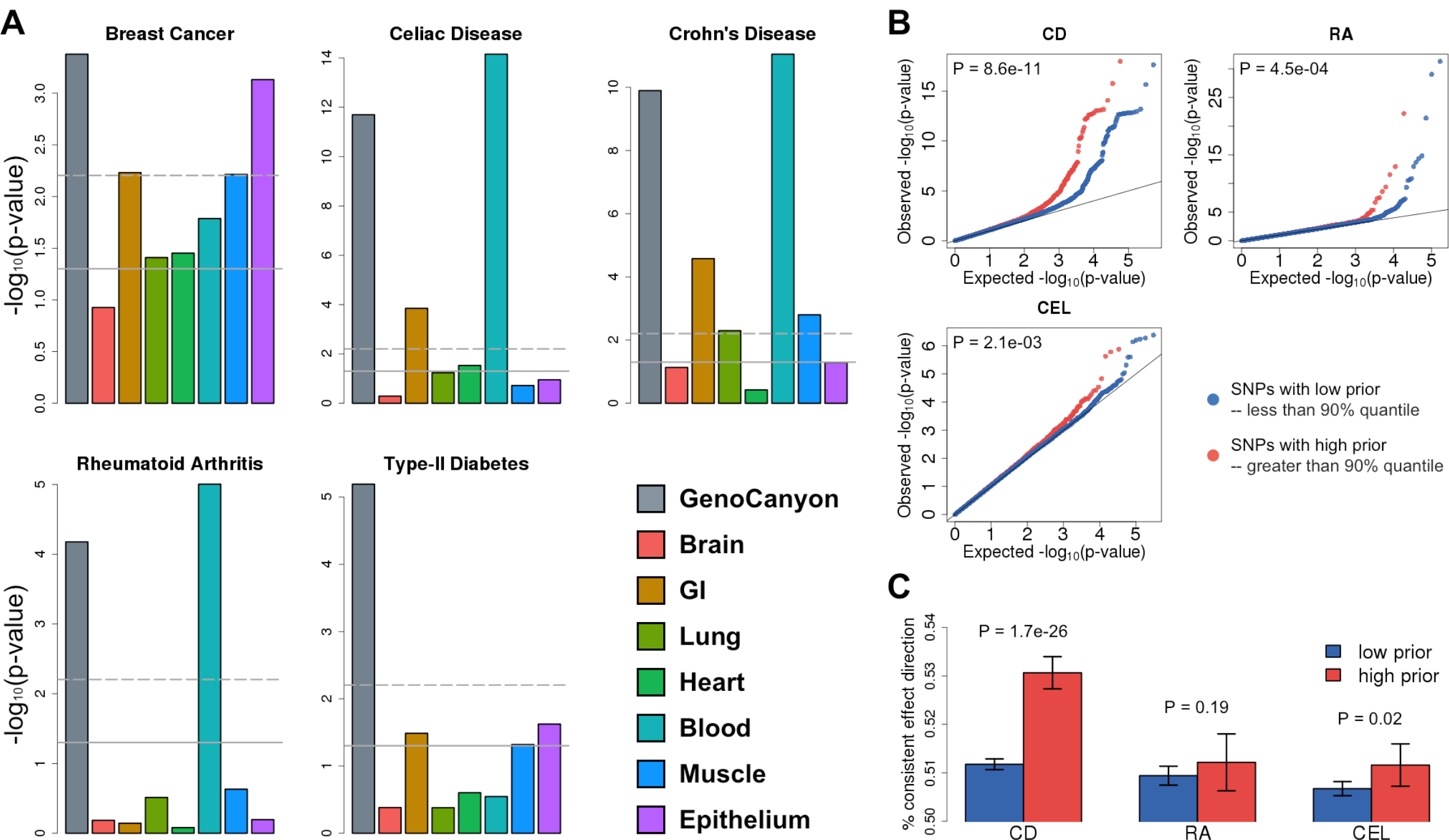
Evaluating effectiveness of annotations and empirical effect size prior. **(A)**GWAS signal enrichment across GenoCanyon and tissue-specific GenoSkyline annotations. The horizontal lines mark p-value cutoffs of 0.05 and Bonferroni corrected significance level. **(B)** Comparing signal strength of SNPs with high priors and low priors in independent validation cohorts. SNPs with higher priors have significantly stronger associations across three independent and well-powered testing datasets (N>2,000). P-values were calculated using one-sided Kolmogorov-Smirnov test. **(C)** Comparing consistency of SNPs’ effect direction between training and testing datasets. Each bar quantifies the proportion of SNPs with consistent effect directions. P-values were calculated using one-sided two-sample binomial test.

Area under the receiver operating characteristic curve (AUC) for different approaches is summarized in **Table 2**. AnnoPred showed consistently improved prediction accuracy compared with all other methods across five diseases. Notably, PRS_sig_ and PRS_all_ showed suboptimal performance in these datasets, reaffirming the importance of modeling LD and other external information. To test different methods’ ability to stratify individuals with high risk, we compared the proportion of cases among testing samples with high PRS. AnnoPred outperformed all other methods in CD, CEL, RA, and T2D (**Supplementary Figure 1**). Next, we tested AnnoPred’s performance using only the 53 baseline annotations and observed a substantial drop in prediction accuracy for all diseases (**Supplementary Table 2**). These results highlight the importance of annotation quality in genetic risk prediction, and also demonstrate GenoCanyon and GenoSkyline’s ability to accurately identify functionality in the human genome.

**Table 2.** AUCs of different methods. The highest AUCs are highlighted in boldface.

| **Disease/Trait** | **PRS<sub>sig</sub>** | **PRS<sub>all</sub>** | **PRS<sub>P+T</sub>** | **LDpred** | **AnnoPred** |
| --- | --- | --- | --- | --- | --- |
| Crohn's Disease | 0.659 | 0.634 | 0.690 | 0.689 | **0.702** |
| Breast Cancer | 0.553 | 0.581 | 0.598 | 0.632 | **0.665** |
| Rheumatoid Arthritis | 0.617 | 0.566 | 0.645 | 0.661 | **0.665** |
| Type-II Diabetes | 0.596 | 0.586 | 0.616 | 0.609 | **0.623** |
| Celiac Disease | 0.576 | 0.593 | 0.624 | 0.631 | **0.640** |

Due to distinct allele frequencies and LD structures across populations, risk prediction accuracy usually drops when the training and testing samples are from different populations. In order to investigate the robustness of AnnoPred against population heterogeneity, we applied AnnoPred to three non-European cohorts for breast cancer and type-II diabetes while training the model using summary statistics from European-based studies. The AUCs are summarized in **Supplementary Table 3**. As expected, we observed a drop in prediction accuracy for all methods. However, AnnoPred still performed the best in all three trans-ethnic validation datasets.

Our work demonstrates that functional annotations can effectively improve performance of genetic risk prediction. AnnoPred jointly analyzes diverse types of annotation data and GWAS summary statistics to provide accurate estimates of SNP effect sizes, which lead to consistently better prediction accuracy for multiple complex diseases. Our method is not without limitation. First, despite the consistent improvement compared with existing PRS-based methods, AUCs for most diseases remain moderate. In order to effectively stratify risk groups for clinical usage, our model remains to be further calibrated using large cohorts with measured environmental and clinical risk factors [1]. Second, accurate estimation of GWAS signal enrichment and SNP effect sizes requires a large sample size for the training dataset. This could be potentially improved by better estimators for annotation-stratified heritability in the future [17]. The rich collection of publicly available integrative annotation data, in conjunction with the increasing accessibility of GWAS summary statistics, makes AnnoPred a customizable and powerful tool. As GWAS sample size continues to grow, AnnoPred has the potential to achieve even better prediction accuracy and become widely adopted as a summary of genetic contribution in clinical applications of risk prediction.

## Methods

### Annotation data

We incorporated GenoCanyon general functionality scores [14], GenoSkyline tissue-specific functionality scores for seven tissue types (brain, gastrointestinal tract, lung, heart, blood, muscle, and epithelium) [15], and 53 LDSC baseline annotations [16] into our model (**Supplementary Table 1**). We smoothened GenoCanyon annotation by taking the mean GenoCanyon score using a 10Kb window as previously suggested [18]. The smoothened GenoCanyon annotation and raw GenoSkyline annotations of seven tissue types were dichotomized based on a cutoff of 0.5. The regions with GenoCanyon or GenoSkyline scores greater than the cutoff were interpreted as non-tissue-specific or tissue-specific functional regions in the human genome. Such dichotomization has been previously shown to be robust against the cutoff choice [15]. Notably, the AnnoPred framework allows users to specify their own choice of annotations.

### Heritability partition

We assume throughout the paper that both the phenotype *Y_N×l_* and the genotypes *X_N×M_* are standardized with mean zero and variance one. We assume a linear model.

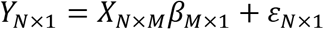

*X*, *β* and *ɛ* are mutually independent. We also assume that *β* is a random effect and effects of different SNPs are independent. A key idea in the AnnoPred framework is to utilize functional annotation information to accurately estimate SNPs’ effect sizes. In order to achieve that, we first partition trait heritability by annotations using LD score regression [16]. Per-SNP heritability is defined as the variance of *β_i_* for the i^th^ SNP, and is used to quantify SNP effect sizes. More specifically, assume there are *K* + 1 predefined annotation categories, denoted as *S*_0_,*S*_1_,…, *S_K_* with *S*_0_ representing the entire genome. Under an additive assumption for heritability in overlapped annotations, we have 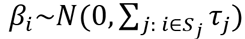, where τ_0_, τ_1_,…, τ_k_ quantify the contribution to per-SNP heritability from each annotation category. Denote the estimated marginal effect size of the i^th^ SNP as 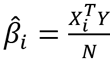, then we have the following approximation

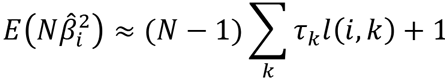

where *l*(*i,k*) is the annotation-stratified LD score and *N* denotes the total sample size. Regression coefficients τ_*k*_ are estimated through weighted least squares. The estimated heritability of the i^th^ SNP is then 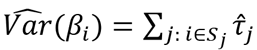

### Empirical prior of effect size

Based on per-SNP heritability estimates, we propose two different priors for SNP effect sizes to add flexibility against different genetic architecture. For the first prior, we assume SNP effect size follows a spike-and-slab distribution

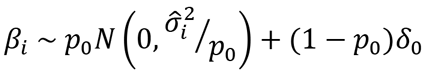

where *p*_0_ is the proportion of causal SNPs in the dataset, and *δ*_0_ is a Dirac function representing a point mass at zero. The empirical variance of each SNP, i.e. 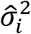, is determined by the annotation categories it falls in. More specifically, we assume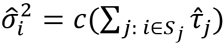, where *c* is a constant calculated from the following equation

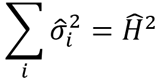

We do not directly use 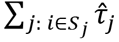 as the empirical variance prior because it is estimated in the context where all SNPs in the 1000 genomes database are included in the model [16]. Such per-SNP heritability estimates cannot be extrapolated to the risk prediction context where much fewer SNPs are analyzed [19]. Therefore, we rescale the heritability estimates to better quantify each SNP’s contribution toward chip heritability. Following [20], we use a summary statistics-based heritability estimator that approximates Haseman-Elston estimator:

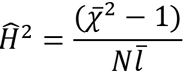

where 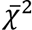 and 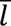 denote mean 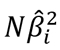 and mean non-stratified LD score, respectively.

In the first prior, we assumed the same proportion of causal SNPs but different effect sizes across annotation categories. We now describe the second prior that assumes different proportions of but the same effect size for causal SNPs. To be specific, we assume causal effect size to be *Var*(*β_causal_*) = *V*, the total number of SNPs to be *M*_0_, and the overall proportion of causal SNPs to be *p*_0_. The total heritability 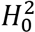 could then be written as 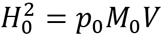. For the i^th^ SNP, use 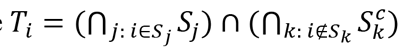 to denote the collection of SNPs that share the same annotation assignment with the i^th^ SNP, and let *M_T_i__* = |*T_i_*|, i.e. number of SNPs in the set. Then, the total heritability of SNPs in *T_i_* is 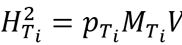, with 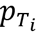 denoting the proportion of causal SNPs in *T_i_*. Following these notations, we have

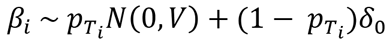

where 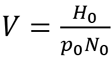 and 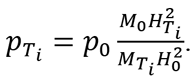. We use 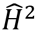 to estimate 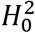, and use the following formula to estimate 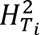.

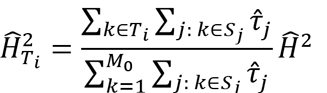

Finally, *p*_0_ is treated as a tuning parameter for both prior functions in our analysis.

### Calculation of posterior effect sizes

By Bayes’ rule, the posterior distribution of *β* is:

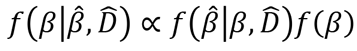

where 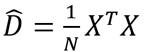 is the sample correlation matrix and 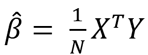 is the marginal effect size estimates. Given *β* and 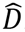, 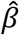 follows a multivariate normal distribution asymptotically with the following mean and variance

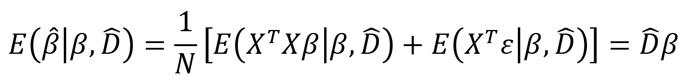

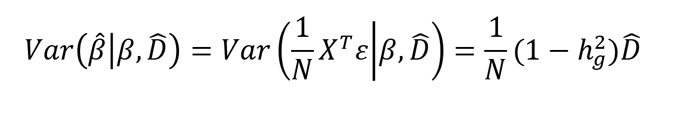

However, 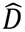 is usually non-invertible and has very high dimensions. We thus study the posterior distribution of a small chunk of 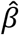 instead. Let 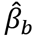 be the estimated marginal effect size of SNPs in a region *b* (e.g. a LD block) and the corresponding genotype matrix is *X_b_* and sample correlation matrix is 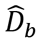. Then the conditional mean and variance of 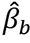 are

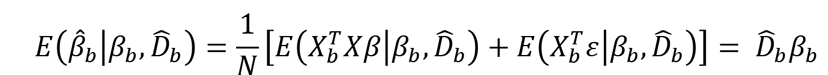

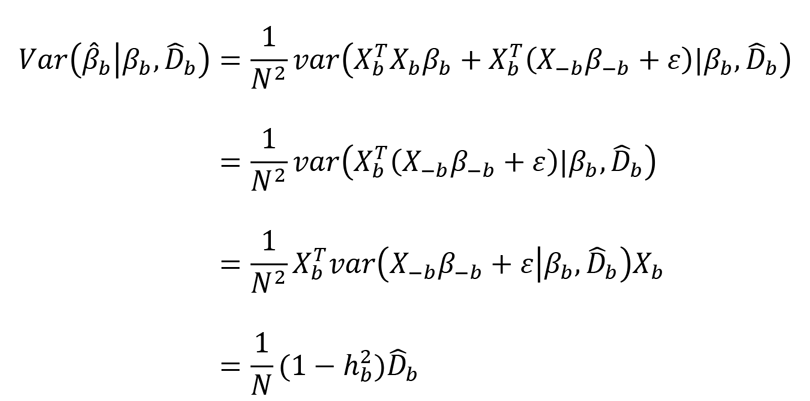

where 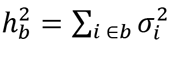 is the heritability of SNPs in region *b*, and *X_−b_* and *β_−b_* denote the genotype matrix and effect sizes of SNPs not in region *b*. The conditional distribution of *β*_*b*_

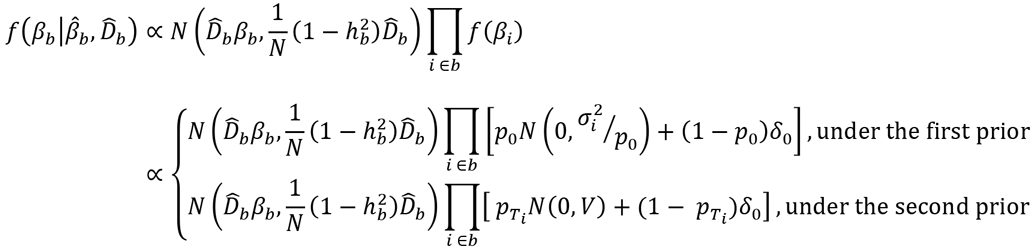

Although it is difficult to derive 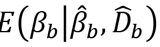 from the joint conditional distribution of β_*b*_, each element of β_*b*_ follows a mixed normal distribution conditioning on 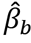, 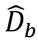, and all other elements in β_*b*_. Therefore, we could apply a Gibbs sampler to draw samples from 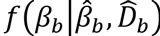, D and use the sample mean as an approximation for 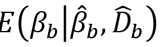.

### Calculation of PRS

PRS is calculated using the following formula

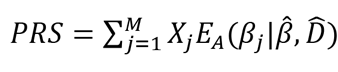

where *E_A_* denotes the posterior expectation as described above. In practice, the individual-level genotype matrix is not available and we use the LD matrix estimated from a reference panel or the validation samples to substitute 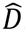. We apply the same standard of choosing the size of *b* as described in [10]. Choices of prior and *p*_0_ can be tuned in an independent cohort. For the data analysis described in this work, we adopted a cross-validation scheme. We tuned parameters using half of the testing samples and evaluated prediction accuracy using the other half, and then repeated the analysis after reversing the two sample subsets. Finally, we reported the mean AUC of two crossvalidations.

### Other methods for comparison

We compared AnnoPred with four commonly used risk prediction methods based on summary data of association studies. PRS_sig_ and PRS_all_ were both calculated as the inner product of marginal effect size estimates and the corresponding genotypes. PRS_all_ used all the SNPs that are shared between training and testing datasets while PRS_sig_ only used SNPs with p-values below 5×10^−8^ in the training set. We downloaded python code for PRS_P+T_ and LDpred from Bitbucket (http://bitbucket.org/bjarni_vilhjalmsson/ldpred).All the tuning parameters were tuned through cross-validation as we did for AnnoPred.

### Simulation settings

We simulated traits from WTCCC genotype data, which contain 15,918 individuals genotyped for 393,273 SNPs after filtering variants with missing rate above 1% and individuals with genetic relatedness above 0.05. We first generated two annotations and each annotation was simulated by randomly selecting 10% of the genome, denoted as *A*_1_ and *A*_2_. Denote the heritability of the trait as 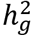 and the number of causal variants as *m* (300 or 3,000). Causal variants were generated as follows:*m*/3 causal variants were selected from *A*_1_, *m*/3 from *A*_2_ and the rest from (*A*_1_⋃*A*_2_)*^C^*. Effect sizes of causal variants were sampled from 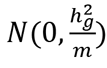. For each simulation, we used 70% of the data to calculate the training summary statistics and randomly divided the rest 30% into two parts for parameter tuning. We also randomly selected half of the training data to calculate summary statistics in order to study the effect of sample size on prediction accuracy.

### GWAS summary statistics and validation data

We trained AnnoPred using publicly accessible GWAS summary statistics and evaluated risk prediction performance using individual-level genotype and phenotype data from cohorts independent from the training samples. Details for each training and testing dataset are provided in **Supplementary Notes** and **Supplementary Table 4**.

For Crohn’s disease, we trained the model using summary statistics from International Inflammatory Bowel Disease Genetics Consortium (IIBDGC; N_case_=6,333 and N_control_=15,056) [21]. Samples from the Wellcome Trust Case Control Consortium (WTCCC) were removed from the meta-analysis and used as the validation dataset (N_case_=1,689 and N_control_=2,891) [22]. For breast cancer, we trained the model using summary statistics from Genetic Associations and Mechanisms in Oncology (GAME-ON) study (N_case_=16,003 and N_control_=41,335) [23], and tested the performance using samples from the Cancer Genetic Markers of Susceptibility (CGEMS) study (N_case_=966 and N_control_=70) [24]. Shared samples between CGEMS and GAME-ON were removed. We used samples from the CIDR-GWAS of breast cancer for trans-ethnic analysis (N_case_=1,666 and N_control_=2,038) [25]. For rheumatoid arthritis, we used summary statistics from a meta-analysis with 5,539 cases and 20,169 controls to train the model [26]. WTCCC samples were removed from the meta-analysis and used for validation (N_case_=1,829 and N_control_=2,892) [22]. For type-II diabetes, the training dataset is Diabetes Genetics Replication and Meta-analysis (DIAGRAM) consortium GWAS with 12,171 cases and 56,862 controls [27]. We used samples from Northwestern NUgene Project for validation (N_case_=662 and N_control_=517) [28]. Samples from Institute for Personalized Medicine (IPM) eMERGE project are used for trans-ethnic analysis (African American:N_case_=517 and N_control_=213; Hispanic:N_case_=477 and N_control_=102) [29]. The training dataset for celiac disease is from a GWAS with 4,533 cases and 10,750 controls [30]. Samples in the National Institute of Diabetes and Digestive and Kidney Diseases (NIDDK) celiac disease study were used for validation (N_case_=1,716 and N_control_=530)[31].

## Software availability

AnnoPred software and source code are freely available online at https://github.com/yiminghu/AnnoPred

## Acknowledgements

This study was supported in part by the National Institutes of Health grants R01 GM59507, the VA Cooperative Studies Program of the Department of Veterans Affairs, Office of Research and Development, and the Yale World Scholars Program sponsored by the China Scholarship Council. We also sincerely thank DIAGRAM, GAME-ON, IIBDGC, and ImmunoBase for making their GWAS summary data publicly accessible. This study makes use of data generated by the Wellcome Trust Case-Control Consortium. A full list of the investigators who contributed to the generation of the data is available from www.wtccc.org.uk. Funding for the project was provided by the Wellcome Trust under award 076113, 085475 and 090355. We also thank Dr. Bjarni J. Vilhjálmsson for sharing his codes.

### Author Contributions

Y.H., Q.L., H.Z. conceived the project and developed the model. Y.H., R.L.P, X.Y. developed the software. Y.H., Q.L., F.F., X.X. performed the analyses. C.Y. contributed collecting and curating data. H.Z. advised on statistical and genetic issues. Y.H., Q.L., and H.Z. wrote the manuscript, and all authors contributed to editing of the manuscript.

### Competing Financial Interests

The authors declare no competing financial interests.

## Supplementary Notes

### Details on GWAS summary statistics and validation data

For Crohn’s disease, we used International Inflammatory Bowel Disease Genetics Consortium (IIBDGC) summary statistics (6,333 Crohn’s disease patients and 15,056 controls) [1]. WTCCC was removed from the meta-analysis and used as a validation set [2]. We filtered individuals with genetic relatedness larger than 0.05 and SNPs with a missing rate larger than 1% and a minor allele frequency less than 1%. In addition, we filtered SNPs with ambiguous nucleotides and kept SNPs matched the summary statistics by both rs number and alleles. After QC, the WTCCC cohort consisted of 1,689 cases and 2,891 controls with 218,833 SNPs overlapping the summary statistics.

For breast cancer, we used the Genetic Associations and Mechanisms in Oncology (GAME-ON) summary statistics, consisting of 16,003 cases and 41,335 controls [3]. As for validation data, we first removed individuals overlapped with BPC3 in GAME-ON from Cancer Genetic Markers of Susceptibility (CGEMS) [4]. The validation set consisted of 966 cases and 70 controls with 497,315 SNPs in common. Besides CGEMS, we also used an African-American as validation data to see how the model performs on different population. The data we used is CIDR-GWAS of Breast Cancer in the African Diaspora (CIDR) [5]. After QC, CIDR consisted of 1,666 cases and 2,038 controls with 555,169 SNPs in common.

For rheumatoid arthritis, we used a meta-analysis consisting of 5,539 cases and 20,169 controls [6]. WTCCC was removed from the meta-analysis and used as a validation set [2]. After QC, WTCCC cohort consisted of 1,829 cases and 2,892 controls with 274,486 SNPs in common.

For type-II diabetes, we used Diabetes Genetics Replication and Meta-analysis (DIAGRAM) consortium GWAS summary statistics with 12,171 cases and 56,862 controls [7]. For testing data, we used Northwestern NUgene Project and after QC it consisted of 662 cases and 517 controls with 479,345 SNPs in common [8].

For celiac disease, we used a GWAS consisting of 4,533 cases and 10,750 controls [9]. The National Institute of Diabetes and Digestive and Kidney Diseases (NIDDK) celiac disease data was used as validation data [10]. After QC, it consisted of 1,716 cases and 530 controls with 504,785 SNPs in common.

### Supplementary Figures

**Supplementary Figure 1.**
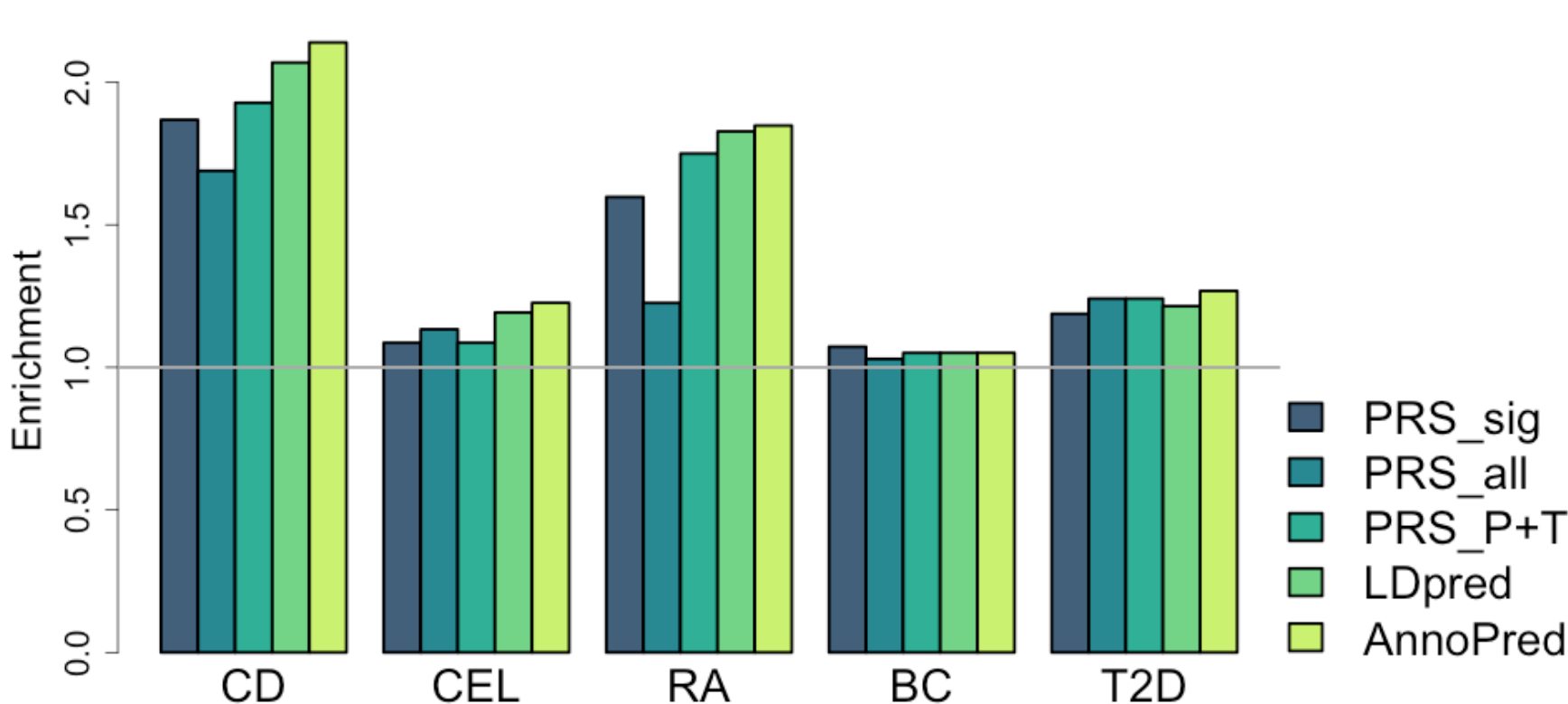
Enrichment of proportion of cases in the top 5% testing samples with high PRS.

### Supplementary Tables

**Supplementary Table 1.** GWAS signal enrichment across 61 annotation categories.

| Category | BC |  |  | CD |  |  | RA |  |  | CEL |  |  | T2D |  |  |
| --- | --- | --- | --- | --- | --- | --- | --- | --- | --- | --- | --- | --- | --- | --- | --- |
|  | Fold | SE | P | Fold | SE | P | Fold | SE | P | Fold | SE | P | Fold | SE | P |
| base_0 | 1 | 0.00 | NA | 1 | 0.00 | NA | 1 | 0.00 | NA | 1 | 0.00 | NA | 1 | 0.00 | NA |
| Coding_UCSC_0 | -0.51 | 6.73 | 8.22E-01 | 8.88 | 5.56 | 1.55E-01 | 2.35 | 10.80 | 9.00E-01 | 6.68 | 4.68 | 2.16E-01 | 2.65 | 6.72 | 8.06E-01 |
| Coding_UCSC.extend.500_0 | 3.57 | 1.73 | 1.40E-01 | 4.77 | 1.10 | 5.62E-04 | 6.24 | 2.20 | 1.48E-02 | 3.41 | 1.44 | 9.28E-02 | 0.50 | 1.76 | 7.77E-01 |
| Conserved_LindbladToh_0 | 15.95 | 6.22 | 8.28E-03 | 4.44 | 3.40 | 3.01E-01 | -5.25 | 6.68 | 3.44E-01 | 10.42 | 5.06 | 5.27E-02 | 4.99 | 5.71 | 4.74E-01 |
| Conserved_LindbladToh.extend.500_0 | 2.17 | 0.62 | 7.47E-02 | 2.46 | 0.35 | 1.10E-04 | 2.90 | 0.60 | 1.54E-03 | 1.76 | 0.46 | 9.87E-02 | 1.63 | 0.58 | 2.74E-01 |
| CTCF_Hoffman_0 | -2.87 | 6.38 | 5.40E-01 | -4.67 | 4.64 | 2.07E-01 | -1.55 | 7.69 | 7.37E-01 | 0.90 | 5.49 | 9.85E-01 | -1.67 | 7.37 | 7.12E-01 |
| CTCF_Hoffman.extend.500_0 | -0.42 | 2.49 | 5.65E-01 | -2.78 | 1.45 | 8.02E-03 | -4.25 | 2.86 | 5.39E-02 | 1.08 | 1.88 | 9.68E-01 | -0.60 | 2.46 | 5.04E-01 |
| DGF_ENCODE_0 | -0.97 | 2.46 | 4.18E-01 | 6.06 | 1.69 | 2.85E-03 | 5.81 | 3.28 | 1.31E-01 | 3.98 | 2.18 | 1.62E-01 | -4.25 | 2.94 | 6.84E-02 |
| DGF_ENCODE.extend.500_0 | 1.35 | 0.43 | 4.27E-01 | 1.87 | 0.25 | 1.67E-03 | 0.81 | 0.56 | 7.21E-01 | 1.29 | 0.34 | 4.01E-01 | 1.18 | 0.42 | 6.75E-01 |
| DHS_peaks_Trynka_0 | 2.82 | 2.81 | 5.15E-01 | 5.86 | 1.75 | 4.05E-03 | 8.19 | 3.80 | 4.42E-02 | 3.03 | 2.50 | 4.17E-01 | -2.71 | 3.59 | 2.91E-01 |
| DHS_Trynka_0 | 4.26 | 2.21 | 1.51E-01 | 2.17 | 1.39 | 3.98E-01 | 2.26 | 2.73 | 6.41E-01 | 2.15 | 1.90 | 5.45E-01 | -0.38 | 2.44 | 5.72E-01 |
| DHS_Trynka.extend.500_0 | 1.17 | 0.50 | 7.26E-01 | 1.40 | 0.33 | 2.24E-01 | 1.07 | 0.65 | 9.11E-01 | 1.06 | 0.44 | 8.88E-01 | 1.46 | 0.49 | 3.60E-01 |
| Enhancer_Andersson_0 | 2.80 | 20.37 | 9.28E-01 | 28.66 | 12.43 | 3.13E-02 | -22.08 | 22.23 | 3.01E-01 | 12.86 | 16.78 | 4.82E-01 | -26.80 | 17.84 | 9.05E-02 |
| Enhancer_Andersson.extend.500_0 | 2.15 | 6.88 | 8.65E-01 | 16.53 | 4.77 | 3.02E-03 | 1.74 | 6.72 | 9.11E-01 | 16.28 | 5.02 | 2.07E-03 | -5.49 | 4.52 | 1.24E-01 |
| Enhancer_Hoffman_0 | 0.68 | 3.70 | 9.30E-01 | 5.36 | 2.03 | 3.06E-02 | 4.07 | 4.28 | 4.65E-01 | 4.48 | 2.61 | 1.76E-01 | -0.03 | 3.26 | 7.53E-01 |
| Enhancer_Hoffman.extend.500_0 | 1.03 | 1.30 | 9.80E-01 | 3.67 | 0.86 | 2.25E-03 | 4.40 | 1.45 | 2.01E-02 | 4.25 | 1.00 | 1.42E-03 | 2.64 | 1.27 | 2.10E-01 |
| FetalDHS_Trynka_0 | -3.21 | 3.63 | 2.12E-01 | 7.21 | 2.15 | 4.87E-03 | 4.04 | 4.38 | 4.75E-01 | 6.92 | 2.81 | 3.08E-02 | -3.65 | 3.84 | 2.15E-01 |
| FetalDHS_Trynka.extend.500_0 | 0.73 | 0.96 | 7.74E-01 | 2.52 | 0.54 | 6.55E-03 | 2.48 | 1.17 | 2.27E-01 | 2.53 | 0.76 | 4.41E-02 | 1.06 | 0.97 | 9.53E-01 |
| H3K27ac_Hnisz_0 | 1.97 | 0.41 | 1.03E-02 | 2.16 | 0.18 | 2.04E-09 | 2.33 | 0.40 | 6.38E-04 | 2.20 | 0.29 | 4.66E-05 | 1.50 | 0.36 | 1.60E-01 |
| H3K27ac_Hnisz.extend.500_0 | 1.97 | 0.33 | 4.03E-03 | 2.11 | 0.21 | 2.01E-07 | 1.84 | 0.44 | 4.66E-02 | 2.26 | 0.28 | 4.32E-06 | 1.92 | 0.33 | 6.58E-03 |
| H3K27ac_PGC2_0 | 0.92 | 0.92 | 9.28E-01 | 2.48 | 0.53 | 4.30E-03 | 2.24 | 1.14 | 2.72E-01 | 3.54 | 0.78 | 6.52E-04 | 1.18 | 0.89 | 8.36E-01 |
| H3K27ac_PGC2.extend.500_0 | 3.20 | 0.55 | 9.59E-05 | 2.51 | 0.32 | 3.25E-06 | 2.85 | 0.73 | 7.75E-03 | 2.52 | 0.45 | 3.98E-04 | 1.84 | 0.53 | 1.20E-01 |
| H3K4me1_peaks_Trynka_0 | 4.01 | 2.14 | 1.61E-01 | 1.92 | 1.37 | 5.04E-01 | -1.53 | 2.42 | 2.92E-01 | 2.98 | 1.65 | 2.14E-01 | 0.20 | 2.16 | 7.07E-01 |
| H3K4me1_Trynka_0 | 1.04 | 0.64 | 9.48E-01 | 1.83 | 0.38 | 3.42E-02 | 0.11 | 0.92 | 3.15E-01 | 2.03 | 0.61 | 8.62E-02 | 1.64 | 0.64 | 3.04E-01 |
| H3K4me1_Trynka.extend.500_0 | 1.74 | 0.23 | 3.52E-03 | 1.62 | 0.15 | 8.41E-05 | 1.86 | 0.28 | 2.39E-03 | 1.72 | 0.20 | 7.31E-04 | 1.57 | 0.24 | 3.54E-02 |
| H3K4me3_peaks_Trynka_0 | -1.61 | 4.67 | 5.80E-01 | 3.68 | 4.28 | 5.35E-01 | 6.85 | 6.64 | 3.71E-01 | 4.70 | 4.86 | 4.44E-01 | -1.90 | 4.59 | 5.26E-01 |
| H3K4me3_Trynka_0 | 2.15 | 1.60 | 4.66E-01 | 4.92 | 1.30 | 2.01E-03 | 2.54 | 2.12 | 4.59E-01 | 2.88 | 1.39 | 1.62E-01 | 1.72 | 1.55 | 6.41E-01 |
| H3K4me3_Trynka.extend.500_0 | 1.55 | 0.68 | 4.19E-01 | 2.59 | 0.53 | 3.03E-03 | 2.29 | 0.94 | 1.72E-01 | 2.30 | 0.59 | 2.68E-02 | 1.78 | 0.70 | 2.70E-01 |
| H3K9ac_peaks_Trynka_0 | 2.10 | 5.70 | 8.45E-01 | 2.74 | 3.22 | 5.94E-01 | 0.50 | 6.41 | 9.38E-01 | 1.43 | 4.62 | 9.25E-01 | -2.29 | 6.15 | 5.90E-01 |
| H3K9ac_Trynka_0 | 3.08 | 2.10 | 2.92E-01 | 3.12 | 1.22 | 8.14E-02 | 3.94 | 2.13 | 1.42E-01 | 1.23 | 1.35 | 8.62E-01 | 3.30 | 1.76 | 1.85E-01 |
| H3K9ac_Trynka.extend.500_0 | 1.16 | 0.90 | 8.55E-01 | 2.25 | 0.52 | 1.75E-02 | 0.18 | 1.08 | 4.31E-01 | 0.89 | 0.75 | 8.81E-01 | 2.58 | 0.80 | 3.98E-02 |
| Intron_UCSC_0 | 1.48 | 0.28 | 6.26E-02 | 1.01 | 0.20 | 9.71E-01 | 1.11 | 0.36 | 7.60E-01 | 1.45 | 0.20 | 1.95E-02 | 1.24 | 0.26 | 3.40E-01 |
| Intron_UCSC.extend.500_0 | 1.49 | 0.23 | 2.10E-02 | 1.28 | 0.17 | 7.83E-02 | 1.18 | 0.22 | 3.97E-01 | 1.52 | 0.19 | 3.54E-03 | 1.14 | 0.18 | 4.26E-01 |
| PromoterFlanking_Hoffman_0 | 8.22 | 10.50 | 4.84E-01 | -9.46 | 6.53 | 1.14E-01 | -15.29 | 14.98 | 2.72E-01 | -7.06 | 8.52 | 3.43E-01 | 9.30 | 11.94 | 4.83E-01 |
| PromoterFlanking_Hoffman.extend.500_0 | 0.84 | 3.73 | 9.66E-01 | 2.35 | 2.44 | 5.80E-01 | 7.41 | 4.45 | 1.46E-01 | 2.83 | 2.84 | 5.16E-01 | 0.77 | 3.99 | 9.54E-01 |
| Promoter_UCSC_0 | 2.47 | 3.28 | 6.53E-01 | 3.85 | 2.39 | 2.36E-01 | 10.22 | 4.39 | 2.82E-02 | 0.99 | 2.57 | 9.96E-01 | 3.84 | 3.57 | 4.20E-01 |
| Promoter_UCSC.extend.500_0 | -0.69 | 1.84 | 3.73E-01 | 4.46 | 1.69 | 3.73E-02 | 5.60 | 2.46 | 5.05E-02 | 2.99 | 1.74 | 2.46E-01 | 1.06 | 1.86 | 9.74E-01 |
| Repressed_Hoffman_0 | -0.65 | 0.64 | 6.29E-03 | 0.06 | 0.35 | 4.36E-03 | 0.42 | 0.66 | 3.82E-01 | 0.11 | 0.51 | 6.82E-02 | 0.39 | 0.64 | 3.34E-01 |
| Repressed_Hoffman.extend.500_0 | 0.81 | 0.16 | 2.04E-01 | 0.51 | 0.09 | 1.29E-07 | 0.48 | 0.20 | 6.04E-03 | 0.37 | 0.16 | 5.75E-06 | 0.74 | 0.16 | 1.07E-01 |
| SuperEnhancer_Hnisz_0 | 2.52 | 0.55 | 4.29E-03 | 2.92 | 0.34 | 7.18E-09 | 2.77 | 0.61 | 4.10E-03 | 4.54 | 0.57 | 1.84E-12 | 1.37 | 0.53 | 4.83E-01 |
| SuperEnhancer_Hnisz.extend.500_0 | 2.75 | 0.49 | 6.27E-04 | 3.03 | 0.34 | 2.41E-09 | 2.57 | 0.53 | 4.23E-03 | 4.45 | 0.53 | 9.33E-14 | 1.41 | 0.48 | 3.94E-01 |
| TFBS_ENCODE_0 | 6.74 | 2.41 | 1.27E-02 | 4.17 | 1.67 | 6.19E-02 | 2.92 | 2.85 | 5.06E-01 | 6.84 | 1.96 | 2.17E-03 | -1.26 | 2.57 | 3.63E-01 |
| TFBS_ENCODE.extend.500_0 | 2.07 | 0.71 | 1.46E-01 | 2.22 | 0.45 | 4.95E-03 | 2.38 | 0.97 | 1.68E-01 | 1.89 | 0.58 | 1.35E-01 | 1.58 | 0.67 | 3.98E-01 |
| Transcribed_Hoffman_0 | 2.24 | 0.80 | 9.63E-02 | 1.17 | 0.46 | 7.09E-01 | 0.86 | 0.81 | 8.62E-01 | 1.16 | 0.57 | 7.74E-01 | 2.19 | 0.78 | 1.02E-01 |
| Transcribed_Hoffman.extend.500_0 | 0.79 | 0.23 | 3.43E-01 | 0.93 | 0.13 | 5.70E-01 | 0.63 | 0.28 | 1.70E-01 | 0.63 | 0.22 | 1.07E-01 | 0.80 | 0.21 | 3.43E-01 |
| TSS_Hoffman_0 | 5.14 | 5.84 | 4.76E-01 | 9.19 | 4.71 | 9.43E-02 | 11.07 | 8.27 | 2.14E-01 | 23.15 | 7.27 | 8.53E-04 | -1.98 | 6.53 | 6.48E-01 |
| TSS_Hoffman.extend.500_0 | 0.32 | 3.42 | 8.42E-01 | 9.21 | 2.41 | 1.31E-03 | 9.75 | 4.45 | 4.37E-02 | 7.56 | 3.26 | 3.49E-02 | -2.42 | 3.56 | 3.31E-01 |
| UTR_3_UCSC_0 | -3.46 | 6.88 | 5.09E-01 | 5.70 | 4.57 | 2.98E-01 | 7.15 | 7.39 | 4.05E-01 | 0.05 | 5.23 | 8.55E-01 | -2.92 | 6.12 | 5.21E-01 |
| UTR_3_UCSC.extend.500_0 | 0.62 | 2.97 | 8.98E-01 | 6.77 | 3.38 | 9.08E-02 | 0.00 | 3.58 | 7.81E-01 | 1.27 | 2.49 | 9.14E-01 | -0.41 | 2.54 | 5.80E-01 |
| UTR_5_UCSC_0 | -8.59 | 10.57 | 3.46E-01 | 3.17 | 8.61 | 8.00E-01 | -6.10 | 18.24 | 6.86E-01 | 11.58 | 8.28 | 1.99E-01 | 15.88 | 24.42 | 5.41E-01 |
| UTR_5_UCSC.extend.500_0 | 3.67 | 3.02 | 3.76E-01 | 3.88 | 2.11 | 1.79E-01 | 6.42 | 3.66 | 1.37E-01 | 1.56 | 2.38 | 8.12E-01 | 2.74 | 3.47 | 6.13E-01 |
| WeakEnhancer_Hoffman_0 | 2.37 | 7.49 | 8.55E-01 | 10.64 | 4.90 | 5.20E-02 | 10.22 | 8.98 | 2.98E-01 | 14.51 | 6.45 | 2.88E-02 | 1.80 | 9.24 | 9.30E-01 |
| WeakEnhancer_Hoffman.extend.500_0 | 2.77 | 1.76 | 3.08E-01 | 2.46 | 1.27 | 2.43E-01 | 4.95 | 2.32 | 1.00E-01 | 3.90 | 1.49 | 6.28E-02 | 3.86 | 1.99 | 1.73E-01 |
| GenoCanyon | 2.04 | 0.29 | 4.20E-04 | 2.37 | 0.19 | 1.27E-10 | 2.56 | 0.41 | 6.66E-05 | 2.76 | 0.29 | 2.01E-12 | 1.92 | 0.26 | 5.82E-04 |
| GenoSkyline-Brain | 3.97 | 2.00 | 1.18E-01 | 3.06 | 1.14 | 7.35E-02 | 1.89 | 2.01 | 6.52E-01 | 1.95 | 1.47 | 5.13E-01 | -0.04 | 1.90 | 5.80E-01 |
| GenoSkyline-GI | 6.61 | 2.11 | 5.87E-03 | 6.03 | 1.17 | 2.64E-05 | 1.66 | 1.82 | 7.18E-01 | 6.25 | 1.52 | 1.41E-04 | 2.25 | 1.65 | 4.40E-01 |
| GenoSkyline-Lung | 9.16 | 3.98 | 3.90E-02 | 6.13 | 1.83 | 5.04E-03 | 4.41 | 3.46 | 3.07E-01 | 5.70 | 2.53 | 5.80E-02 | 0.76 | 3.31 | 9.42E-01 |
| GenoSkyline-Heart | 5.99 | 2.51 | 3.53E-02 | 2.34 | 1.49 | 3.77E-01 | 1.58 | 2.72 | 8.30E-01 | 5.48 | 2.15 | 2.92E-02 | -0.86 | 2.56 | 4.66E-01 |
| GenoSkyline-Blood | 5.52 | 1.96 | 1.63E-02 | 9.59 | 1.19 | 8.93E-12 | 10.00 | 2.41 | 9.89E-06 | 13.92 | 2.03 | 7.03E-15 | 1.87 | 1.61 | 5.84E-01 |
| GenoSkyline-Muscle | 7.37 | 2.50 | 6.12E-03 | 5.47 | 1.38 | 1.59E-03 | 3.69 | 2.27 | 2.34E-01 | 3.33 | 1.83 | 1.91E-01 | 3.92 | 2.28 | 1.83E-01 |
| GenoSkyline-Epithelium | 7.56 | 2.10 | 7.40E-04 | 3.95 | 1.49 | 5.15E-02 | 2.06 | 2.31 | 6.38E-01 | 3.45 | 1.64 | 1.11E-01 | 2.88 | 1.67 | 2.48E-01 |

**Supplementary Table 2.** Comparison of the complete model and AnnoPred with baseline annotations. The highest AUCs are highlighted in boldface.

| **Disease/Trait** | **AnnoPred<sub>baseline</sub>** | **AnnoPred<sub>complete</sub>** |
| --- | --- | --- |
| Crohn's Disease | 0.673 | **0.702** |
| Breast Cancer | 0.552 | **0.665** |
| Rheumatoid Arthritis | 0.536 | **0.665** |
| Type-II Diabetes | 0.587 | **0.623** |
| Celiac Disease | 0.608 | **0.640** |

**Supplementary Table 3.**
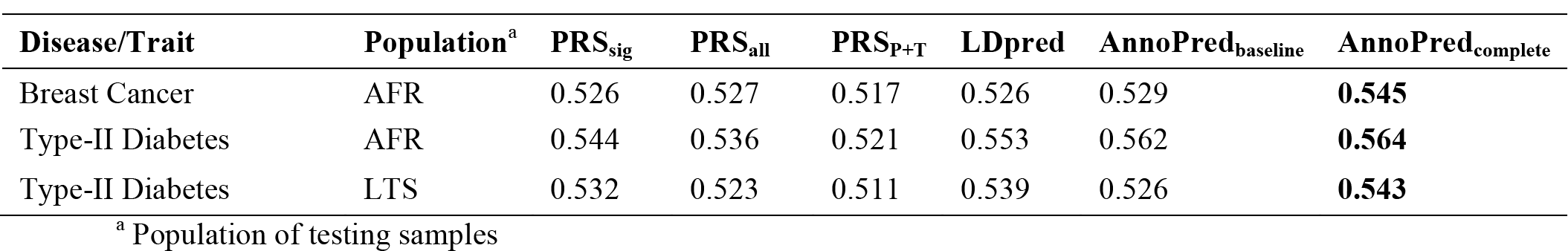
AUCs for trans-ethnic analyses. The highest AUCs are highlighted in boldface.

| Disease/Trait | Population<sup>a</sup> | PRS<sub>sig</sub> | PRS<sub>all</sub> | PRS<sub>P+T</sub> | LDpred | AnnoPred<sub>baseline</sub> | AnnoPred<sub>complete</sub> |
| --- | --- | --- | --- | --- | --- | --- | --- |
| Breast Cancer | AFR | 0.526 | 0.527 | 0.517 | 0.526 | 0.529 | **0.545** |
| Type-II Diabetes | AFR | 0.544 | 0.536 | 0.521 | 0.553 | 0.562 | **0.564** |
| Type-II Diabetes | LTS | 0.532 | 0.523 | 0.511 | 0.539 | 0.526 | **0.543** |

<sup>a</sup> Population of testing samples

**Supplementary Table 4.** URLs for training and testing datasets.

| **Disease/Trait** | **GWAS summary statistics** | **Validation datasets** |
| --- | --- | --- |
| Crohn's Disease | http://www.ibdgenetics.org | http://www.wtccc.org.uk/ccc1/wtccc1_studies.html |
| Breast Cancer | http://gameon.dfci.harvard.edu | http://www.ncbi.nlm.nih.gov/projects/gap/cgi-bin/study.cgi?study_id=phs000147.v3.p1<br>http://www.ncbi.nlm.nih.gov/projects/gap/cgi-bin/study.cgi?study_id=phs000383.v1.p1 |
| Rheumatoid Arthritis | http://www.broadinstitute.org/ftp/pub/rheumatoid_arthritis/Stahl_etal_2010NG/ | http://www.wtccc.org.uk/ccc1/wtccc1_studies.html |
| Type-II Diabetes | http://diagram-consortium.org/downloads.html | http://www.ncbi.nlm.nih.gov/projects/gap/cgi-bin/study.cgi?study_id=phs000237.v1.p1 |
| Celiac Disease | https://www.immunobase.org/downloads/protected_data/GWAS_Data/ | http://www.ncbi.nlm.nih.gov/projects/gap/cgi-bin/study.cgi?study_id=phs000274.v1.p1 |

